# Probabilistic mapping of human functional brain networks identifies regions of high group consensus

**DOI:** 10.1101/2020.09.28.313791

**Authors:** Ally Dworetsky, Benjamin A. Seitzman, Babatunde Adeyemo, Maital Neta, Rebecca S. Coalson, Steven E. Petersen, Caterina Gratton

## Abstract

Many recent developments surrounding the functional network organization of the human brain have focused on data that have been averaged across groups of individuals. While such group-level approaches have shed considerable light on the brain’s large-scale distributed systems, they conceal individual differences in network organization, which recent work has demonstrated to be common and widespread. This individual variability produces noise in group analyses, which may average together regions that are part of different functional systems across participants, limiting interpretability. However, cost and feasibility constraints may limit the possibility for individual-level mapping within studies. Here our goal was to leverage information about individual-level brain organization to probabilistically map common functional systems and identify locations of high inter-subject consensus for use in future group analyses. We probabilistically mapped 14 functional networks in multiple datasets with relatively high amounts of data. All networks show “core” (high-probability) regions, but differ from one another in the extent of their higher-variability components. These patterns replicate well across four datasets with different participants and scanning parameters. We produced a set of high-probability regions of interest (ROIs) from these probabilistic maps; these and the probabilistic maps are made publicly available, together with a tool for querying the network membership probabilities associated with any given cortical location. These quantitative estimates and public tools may allow researchers to apply information about inter-subject consensus to their own fMRI studies, improving inferences about systems and their functional specializations.

## INTRODUCTION

A key objective of functional magnetic resonance imaging (fMRI) studies has been to gain insight into how brain regions respond during tasks and how they interact with one another in distributed large-scale systems. To do so, analyses have typically been performed on averages across groups of subjects, to counteract noisy data from individuals. Studies using a group-average approach to examine human functional brain networks have produced robust and well-validated descriptions of, for example, typical functional network architecture (Power et al., 2011; Yeo et al., 2011).

Although the group-average approach has been useful in revealing fundamental qualities of functional network organization, recent data have suggested that averaging across subjects ignores distinct individual-specific features of cortical organization (Braga & Buckner, 2017; Finn et al., 2015; Gordon et al., 2017a; Kong et al., 2019; Miranda-Dominguez et al., 2014; Mueller et al., 2013). Historically, a major barrier to producing reliable connectivity estimates at the individual level using resting-state functional connectivity (RSFC) techniques has been acquiring a sufficient quantity of data to counteract sampling variability (Gordon et al., 2017c; Laumann et al., 2015). Previous work has demonstrated that the reproducibility of connectivity estimates and individual-specific features of functional brain networks is drastically improved with greater quantities of data per subject (Anderson et al., 2011; Elliott et al., 2019; Laumann et al., 2015; Noble et al., 2017) Accordingly, RSFC studies acquiring a typical 5-10 minutes of data per subject may not be sufficient to accurately reflect connectivity patterns in a given individual, or to examine individual differences in network organization. Several recent works have used higher reliability datasets to illuminate regions of high individual differences in functional network topography (Braga & Buckner, 2017; Gordon et al., 2017a; Seitzman et al., 2019), outlining a geography of brain locations that show substantial variability across individuals.

Given known individual differences in functional networks, experimenters are posed with the dilemma of how to continue with analyses of these systems in their own work. One possibility is to acquire sufficient “precision” fMRI data to overcome sampling variability and produce accurate measures of individual brain networks. However, this may be expensive, difficult in certain participant groups, and not possible in previously acquired datasets. An alternative is to quantify the degree of consensus in network profiles across individuals and focus analyses on locations with known commonalities. Despite individual differences, past data have suggested that commonalities in network organization are also large and widespread, with many regions of the cortex showing substantial similarity to the typical group-average brain (Gratton et al., 2018; Kong et al., 2019; Seitzman et al., 2019). The locations of consensus in functional networks can be derived by probabilistically mapping networks across individuals where sufficient fMRI data is available to achieve good individualized network estimates. Consensus locations from these probabilistic maps can then be used to enhance group analyses by (1) reducing heterogeneity (due to averaging different systems across participants), (2) limiting confounds from mixing diverse systems across individuals (e.g., allowing researchers to better understand functional specialization of different brain systems), and (3) determining the extent to which group data can be extrapolated to single subjects. For instance, one could use these maps to determine if an elicited activity pattern maps on to the frontoparietal network or a combination of networks across subjects.

In the present work, we aimed to address this need by probabilistically mapping functional networks across participants in four different datasets. With this information, we can quantify areas of high group consensus: regions where the greatest group convergence in functional network organization is observed across individuals. We provide tools that can be directly applied in various experimental contexts to quantify the degree of consistency in network assignments across a group. The quantitative probabilistic description of functional networks as well as the tools for implementing high-consensus group analyses are likely to be useful to many in the field with insufficient data to map individualized brain networks.

To create high quality estimates of group consensus, we focused our analyses on datasets with relatively high amounts of resting-state data per person (“highly-sampled datasets” > 20 min. of low-motion resting-state data), where individual network maps achieve higher reliability. We used a template-matching procedure to identify cortical brain networks in these highly-sampled individuals and combined the resulting maps to produce a cortex-wide probabilistic estimate for each network. We replicated these findings across four datasets (a Dartmouth dataset with N = 69 with > 20 min. of data per person as the primary dataset, and secondary replications in the Midnight Scan Club: N = 9 with > 154 min., the Human Connectome Project: N = 384, with > 43 min., and a Yale dataset: N = 65, with > 22 min.). Notably, each of these datasets were conducted both on different individuals and with varied scanning parameters. Probabilistic maps are presented and quantified at various thresholds and are validated by contrasting to past independent results of high variability regions). Finally, we provide two tools for research use: (1) a set of network-specific, high-probability ROIs for use in seeding group analyses and (2) a point-and-click tool allowing researchers to explore voxel-by-voxel probabilistic network estimates for regions of activation in their own data. The use of high-consensus regions may provide greater confidence in ROIs selected as priors in network-informed resting-state studies, with the potential for use in task-based studies as well.

## METHODS

### Datasets and overview

Five independent datasets focused on young neurotypical populations were utilized in this paper (Table 1): a Washington University dataset (a subset of the participants reported in Power et al., 2012), a Dartmouth dataset (Gordon et al., 2016), the Midnight Scan Club (MSC) dataset (Gordon et al., 2017c), the Human Connectome Project (HCP) dataset (Van Essen et al., 2012b), and the Yale Low-res dataset (Scheinost et al., 2016; note this dataset extended to middle age). Each dataset we use here consists of highly sampled subjects with a relatively large amount of low-motion data, ranging from a minimum of 20 min. (for N = 69 in the Dartmouth primary mapping dataset) to upwards of 154 min. (for N = 9 in the MSC replication dataset). This large amount of data dramatically increases the reliability of functional connectivity measurements relative to more typical 5-10 min. scans (Gordon et al., 2017c; Laumann et al., 2015).

**Table 1.**
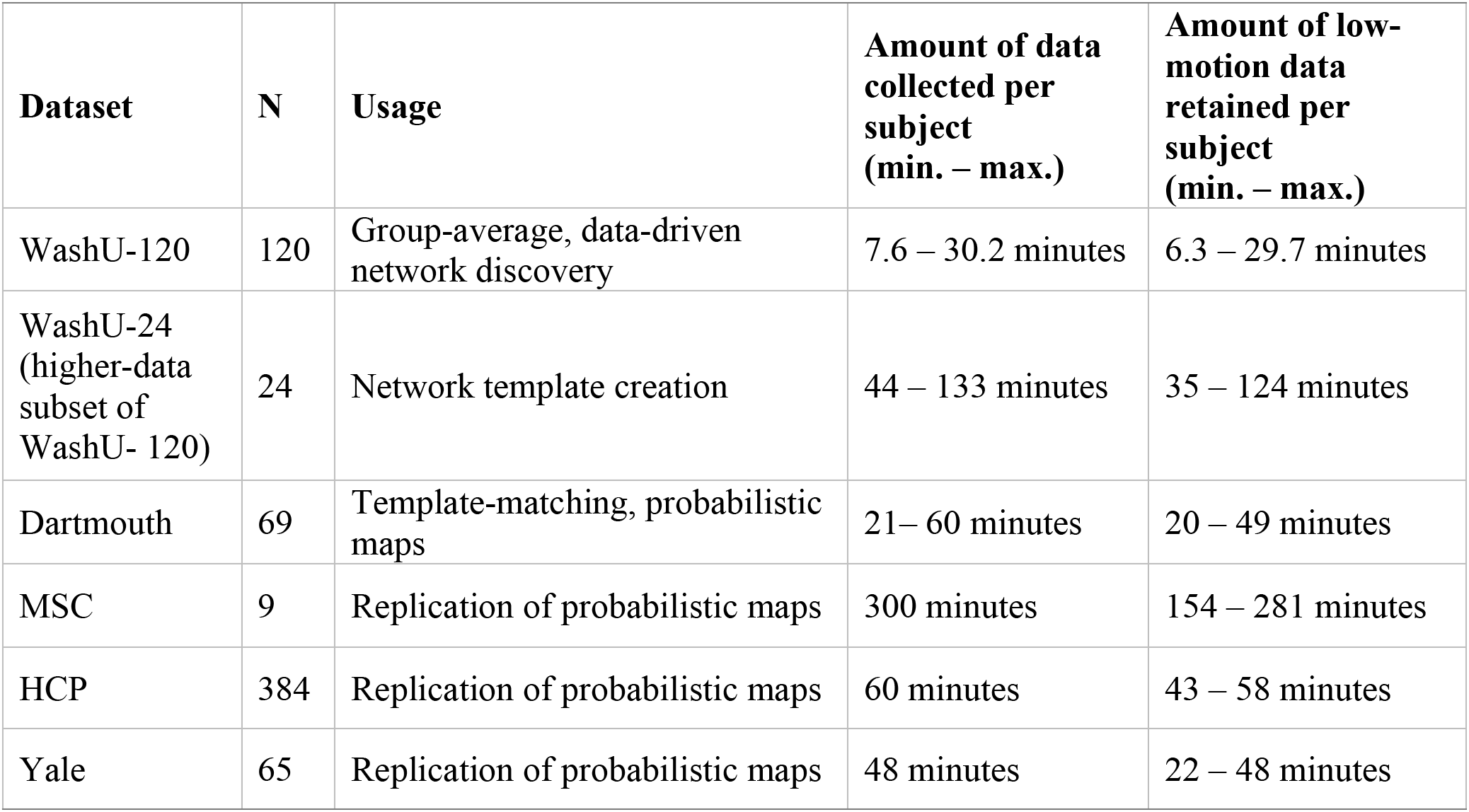
Datasets and data details. Low-motion data quantities were measured after correction for movements using framewise displacement (see *Functional connectivity processing*).

The WashU datasets were used to generate network templates: first using the WashU-120 (60 female, average age 24.7 years) was used to create a data-driven group-average cortical network classification and then a subject “subset” of the WashU-120 consisting of 24 highly sampled subjects (the “WashU-24”) was used to create a set of high-quality average templates based on these group-average networks. Subjects in this subset had at least 35 min. of low-motion data when combining across additional resting-state scan sessions previously obtained from our group (see *Template Generation* in the supplement for more details).

These group-average templates were then applied to subjects in the Dartmouth dataset to identify brain networks in single individuals. The Dartmouth dataset (N = 69 subjects [56 female; average age 20.2 years]) included subjects with over 20 min. of low-motion data. Given its relatively large sample size and its standard, single-band scanning parameters, this dataset was the primary dataset used to determine network probabilities across individuals and generate network-specific regions of high inter-subject consensus.

Three additional datasets were used to replicate these probabilistic maps: the MSC dataset (N = 9 subjects [4 female; average age 29.3 years] with over 154 min. of low-motion rest data), subjects from the HCP dataset (N = 384 subjects [210 female; average age 28.4 years] with at least 52 minutes of data), and subjects from the Yale dataset (N = 65 subjects [32 female; average age 32.2 years; subject ages in this dataset ranged higher] with over 22 minutes of data). Notably, the MSC dataset includes very highly sampled individuals whose functional connectivity maps have been demonstrated to have high reliability and validated with functional activation studies. The HCP dataset replicates the current findings in a large dataset at high spatial and temporal resolution, and the Yale data set replicates the findings in a relatively “low-resolution” dataset (voxel size 3.4 × 3.4 × 6mm). See Supp. Table 1 for acquisition parameters for functional data across all datasets; details on all preprocessing and functional connectivity (FC) processing procedures are outlined below.

#### Preprocessing and FC processing of BOLD data

##### WashU, Dartmouth, MSC, Yale datasets

All structural and functional data were preprocessed to remove noise and artifacts, following Miezin et al. (2000).

###### Structural and functional preprocessing

In the WashU, Dartmouth, MSC, and Yale datasets, slice timing correction was performed using sinc interpolation to account for temporal misalignment in slice acquisition time. Next, whole-brain intensity values across each BOLD run were normalized to achieve a mode value of 1000. Motion correction was performed within and across BOLD runs via a rigid body transformation. Functional BOLD data was then registered either directly to a high resolution T1-weighted structural image from each participant (WashU, Dartmouth, Yale, and HCP datasets) or first to a T2-weighted image and then to the T1 (MSC) using an affine transformation. This T1-weigthed image was aligned to a template atlas (Lancaster et al., 1995) conforming to Talairach stereotactic atlas space (Talairach & Tournoux, 1988) using an affine transformation. All computed transformations and resampling to 3 mm isotropic voxels were simultaneously applied at the end of these steps. For some supplemental analyses to test the effects of structural alignment procedures, cortical surfaces were also generated by FreeSurfer (Dale et al., 1999), registered to fs_LR surface space (Van Essen et al., 2012a), and aligned with each individual’s functional data using the processing stream described in Gordon et al. (2016).

###### Functional connectivity processing

Following Power et al. (2014), additional denoising was applied to the resting-state data for FC analysis. Temporal masks for each subject’s timeseries were created in the WashU, MSC, and Yale datasets by censoring all frames with a framewise displacement (FD; Power et al., 2012) greater than 0.2 mm, and in the Dartmouth dataset by censoring frames with FD greater than 0.25 mm. This frame-censoring approach was implemented to remove timepoints associated with motion, as even small movements can induce distance-dependent biases in functional connectivity (Power et al., 2014, 2018; Satterthwaite et al., 2019), and censoring of high-motion frames has been shown to be effective in reducing distance-dependent artifacts (Ciric et al., 2017, 2018). Across all datasets, segments with fewer than 5 contiguous frames were censored. FreeSurfer 5.0 segmentation using each subject’s T1 image generated a white matter and a cerebrospinal fluid nuisance mask per individual. After BOLD data was demeaned and detrended, Regression of nuisance signals was implemented, regressing out global signal, cerebrospinal fluid, and white matter, as well as the six rigid-body motion regressors and their expansion terms (Friston et al., 1996). Data from high-motion frames were interpolated over via a spectra-matching interpolation technique. Data were then bandpass temporally filtered between 0.009 Hz to 0.08 Hz. Finally, the data were spatially smoothed at FWHM (6 mm).

##### HCP dataset

Preprocessing and FC processing of HCP subjects were carried out similarly to the other datasets with a few differences. First, slice-timing correction was not performed, following the recommendations of the minimal preprocessing pipeline guidelines (Glasser et al., 2013). Second, prior to censoring high-motion frames, motion parameters were low-pass filtered at 0.1 Hz to mitigate effects of respiratory artifacts on motion estimates attributable largely to the multi-band, fast-TR data acquisition (Fair et al., 2020; Siegel et al., 2017). Following this, a filtered FD threshold of 0.1 mm was applied to censor frames. Data were originally processed in MNI atlas space with 2 mm isotropic voxels and were transformed into Talairach space with 3 mm isotropic voxels in a single step prior to spatial smoothing as described above.

#### Template-matching and generation of high-probability ROIs

In this work, we created network maps for highly sampled individual subjects using a template matching approach. These network maps were then overlaid to generate a probabilistic estimate of network distributions across subjects. High-consensus ROIs were generated for research use from regions of high cross-subject agreement of network assignment. Procedures for template-matching in individuals and probabilistic network map generation are illustrated in Fig. 1 and described in more detail below. All analyses were performed in volume (Talairach) space with 3 mm isotropic voxels (figures in this manuscript show data projected to the cortical surface for visualization purposes only).

**Figure 1:**
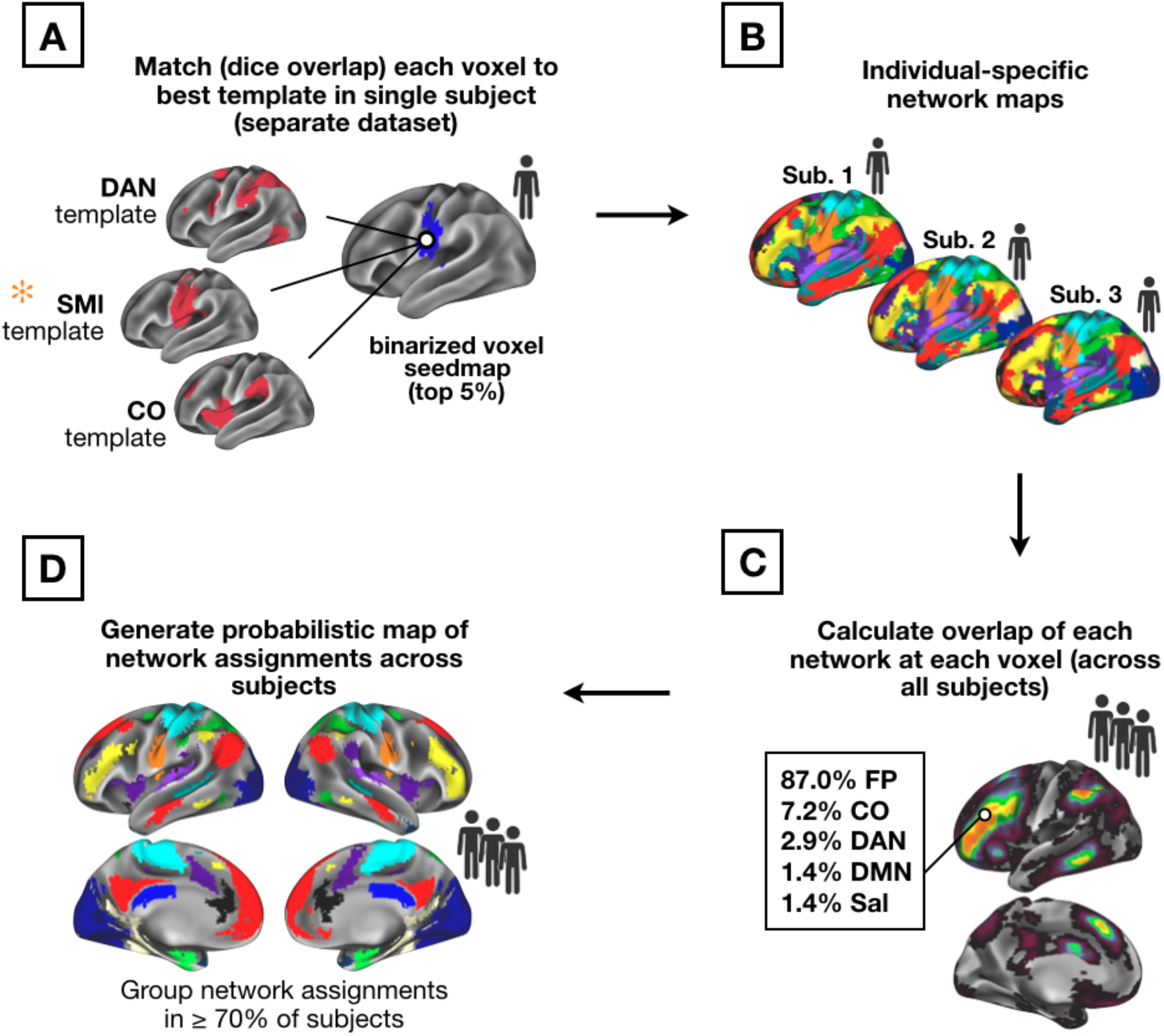
Template-matching procedure (*A)* and creation of probabilistic network maps (*B-D*). A set of group-average network templates were created from the WashU dataset. These group-average templates were binarized at the top 5% of connectivity values. Next, for each single individual in the Dartmouth and replication datasets, a voxelwise correlation matrix was calculated between all gray matter voxels. Seedmaps were thresholded at the top 5% of values across voxels. The individual’s voxel-level binarized map was then iteratively compared (by Dice overlap) with each group-average network template (also thresholded to the top 5% of values), and the network with the highest Dice coefficient was assigned to the voxel (*A*). Once all voxels were assigned in all subjects (*B*), the number of network assignments at each voxel were tallied across subjects (*C*) to generate probabilistic maps of networks. These probabilistic maps were then thresholded (*D*) to represent locations with network consensus in a large majority of subjects. Note that while all steps were performed in volume (Talairach) space, results are mapped onto a template surface for visualization purposes only.

##### 1. Template-matching

Brain networks were identified in individual subjects by a winner-take-all procedure (similar to that employed in Gordon et al. (2017b) which assigned each cortical gray matter voxel in a particular subject to one of 14 network templates. The generation of volumetric network templates is described in the *Supplemental Methods*. Networks include the default mode (DMN), visual, fronto-parietal (FP), dorsal attention (DAN), language (Lang.; this corresponds with the network labeled as “ventral attention” in previous work from our group), salience, cingulo-opercular (CO), somatomotor dorsal (SMd), somatomotor lateral (SMl), auditory, temporal pole (Tpole), medial temporal lobe (MTL), parietal medial (PMN), and parieto-occipital (PON; sometimes called the retrosplenial, contextual association, or parahippocampal systems). Note that the functional/anatomical nomenclature associated with network labels is a matter of ongoing debate (Uddin et al., 2019; here we selected names consistent with previous iterations from Laumann et al., 2015 and Power et al., 2011).

Networks were matched in each individual by assigning each voxel to one of the 14 canonical networks based on the voxel seedmap’s “fit” with each network template (similar to the approach implemented in past work; e.g., Gordon et al., 2017b). Specifically, a seedmap was created for each location and binarized to the top 5% of connectivity values across voxels (this threshold was set based on previous work, but Gordon et al. (2017a) demonstrated consistent network assignments within a subject across a range of individual-level connectivity thresholds). Each voxel’s binarized map was iteratively compared with the 14 network templates (also binarized, see Supp. Fig. 1) and matched to its “best fit.” Fit was measured using the Dice coefficient of overlap between the binarized voxel connectivity map and each binarized template map (Fig. 1A). This procedure was repeated across all cortical voxels, resulting in a cortex-wide individual-specific network map (Fig. 1B). Same-network clusters of less than 108 mm3 (4 contiguous voxels) were removed from each individual’s network map.

Rather than using a data-driven community detection approach to map individualized networks, this template-matching approach was chosen based on our goal of investigating known, previously described brain networks to allow for a reliable comparison of network structure across individuals. However, supplemental analyses in the Midnight Scan Club dataset show comparisons between probabilistic maps derived from data-driven vs. template-based assignments (e.g., see Supp. Fig. 5).

##### 2. Creating probabilistic maps

After individual-specific network maps from the Dartmouth dataset had been generated with the template mapping procedure, these maps were overlapped to produce a cross-subject probabilistic map for each network (Fig. 1C). To generate this cross-subject probabilistic map, individual network assignments at each brain location were tallied to calculate the total occurrence (in number of subjects, with a given network assignment out of the total N = 69). This produced a continuous probabilistic map for each network which specified the probability of a given network assignment at every voxel within the cortical mask. Frequency values of network assignments were divided by the number of subjects within the primary dataset and were converted to percentages to illustrate the probability of network membership at each voxel. Probabilistic maps were created in the same manner from the MSC, Yale, and HCP datasets based on the number of subjects included (9, 65, and 384, respectively), and were compared to the results from the primary dataset. Thresholded versions of the network-specific probabilistic maps were also produced (Fig. 1D), allowing for visualization of the network assignment frequencies at various probability thresholds (e.g., in 50, 60, 70, 80, or 90 percent of subjects). Network-specific probabilistic maps for the Dartmouth and HCP datasets were published at https://github.com/GrattonLab.

Two approaches were taken to quantify the similarity of probabilistic network maps between the primary and replication datasets. First, we calculated the spatial correlation between the unthresholded probability maps across each dataset for each network. Second, we conducted a network-wise random rotation analysis on the thresholded high-consensus locations similar to Gordon et al. (2016). Each network in the (volume-to-surface-mapped) 70% probability map from the Dartmouth dataset was randomly rotated around the 32k_fs_LR cortical surface such that it maintained its size and shape. This rotation was repeated 1000 times for each network in each hemisphere. For each of rotation, we calculated the Dice coefficient between the randomly rotated network in the Dartmouth dataset and the thresholded 70% probability map in each of the replication datasets (MSC, Yale, and HCP). Iterations where a network rotated into the medial wall were ignored and these Dice values were assigned with the average coefficient across all random rotations for that network (Gordon et al., 2016). The similarity between the original (true) Dartmouth consensus map and the replication maps was also assessed via a Dice coefficient. Finally, a p-value was calculated based on the proportion of rotations in which the rotated Dice value exceeded the true Dice value.

##### 3. Creating ROIs of high group consistency for studies in other modalities

Once probabilistic maps were defined, we next set out to create a set of regions of interest (ROIs) with high group consensus for use in future (and retrospective) studies. These ROIs were created by contrasting the probabilistic maps generated above from the Dartmouth dataset with 248 (of 264) ROIs of the larger set previously proposed in Power et al. (2011) found in the cerebral cortex.

Specifically, high group consensus regions were derived from the probabilistic maps of the Dartmouth dataset by identifying locations that showed consistent network assignments across a large majority (i.e., > 75%) of subjects. A spherical 7 mm diameter region was placed on each of the center coordinates reported in Power et al. (2011). ROIs were identified as “high-probability” if their average probability (across voxels) was ≥ 75%. If a region failed to meet the 75% criteria to be identified as “high-probability,” it was shifted one voxel in space (i.e., 3 mm in the x, y, or z direction) and was retained if this shift produced an average ROI probability that met the threshold, as the intention was to keep the original ROI set relatively intact but optimized for probabilistic mapping. As a result of this procedure, a total of 44 ROIs were shifted from their original position. ROIs that failed to meet the high consensus definition with a single voxel shift were dropped from the final group. The full probabilistic maps are also provided to the public, allowing authors the possibility to generate additional ROIs with more varied characteristics if desired.

##### 4. Creating point and click voxel-wise network tool

Finally, we created a tool for displaying the probability of network membership at each cortical voxel for research use. Specifically, a scene file was created using the Connectome Workbench software that contains each network’s probabilistic map in volume space and allows for point-and-click usability to identify the probability (across subjects) that a given voxel is associated with each network.

##### 5. Data and code availability

High consensus ROIs and full probabilistic maps are provided at https://github.com/GrattonLab (ROIs are provided as center coordinates so they can be shaped to a preferred size; coordinates are provided in both Talairach and MNI space for convenience). Data from the Midnight Scan Club is available at https://openneuro.org/datasets/ds000224; data from the Human Connectome Project can be accessed at https://db.humanconnectome.org/; data associated with the WashU-120 is available at https://openneuro.org/datasets/ds000243/versions/00001.

## RESULTS

### Overview of results

In this work, we sought to characterize high-consensus network locations for use in analysis and interpretation of group research studies. To this end, we compiled individualized network assignments in several datasets of highly sampled subjects to create a reliable cross-subject probabilistic map of network definitions. We show these results first for the Dartmouth dataset (primary) and then replicate these findings in three additional datasets to demonstrate their stability. Using these datasets, we explored the degree of consensus for networks across different probability thresholds. Finally, we created two tools for use in future research studies: (1) a set of “high-probability” regions of interest, and (2) a publicly available point-and-click tool for determining network probabilities in researcher-specified locations.

#### Estimated probabilistic maps of 14 canonical networks

As described in the Methods, we used a template-matching approach to determine a voxel-based network assignment for each individual (N = 69) in our primary Dartmouth dataset, based on templates created from the WashU cohort. We then overlapped the individuals’ network maps within the Dartmouth dataset for each canonical network. This overlap was used to generate a cross-subject probabilistic map (Fig. 2; see Supp. Fig. 2 for maps for the remaining 8 canonical networks we examined). As can be seen, all networks demonstrated some regions with high-group consensus (warm colors), but also a spread of lower-consensus locations. While this analysis was conducted in volume space, analyses performed with surface-based alignment produced similar results in the primary dataset (Supp. Fig. 3).

**Figure 2:**
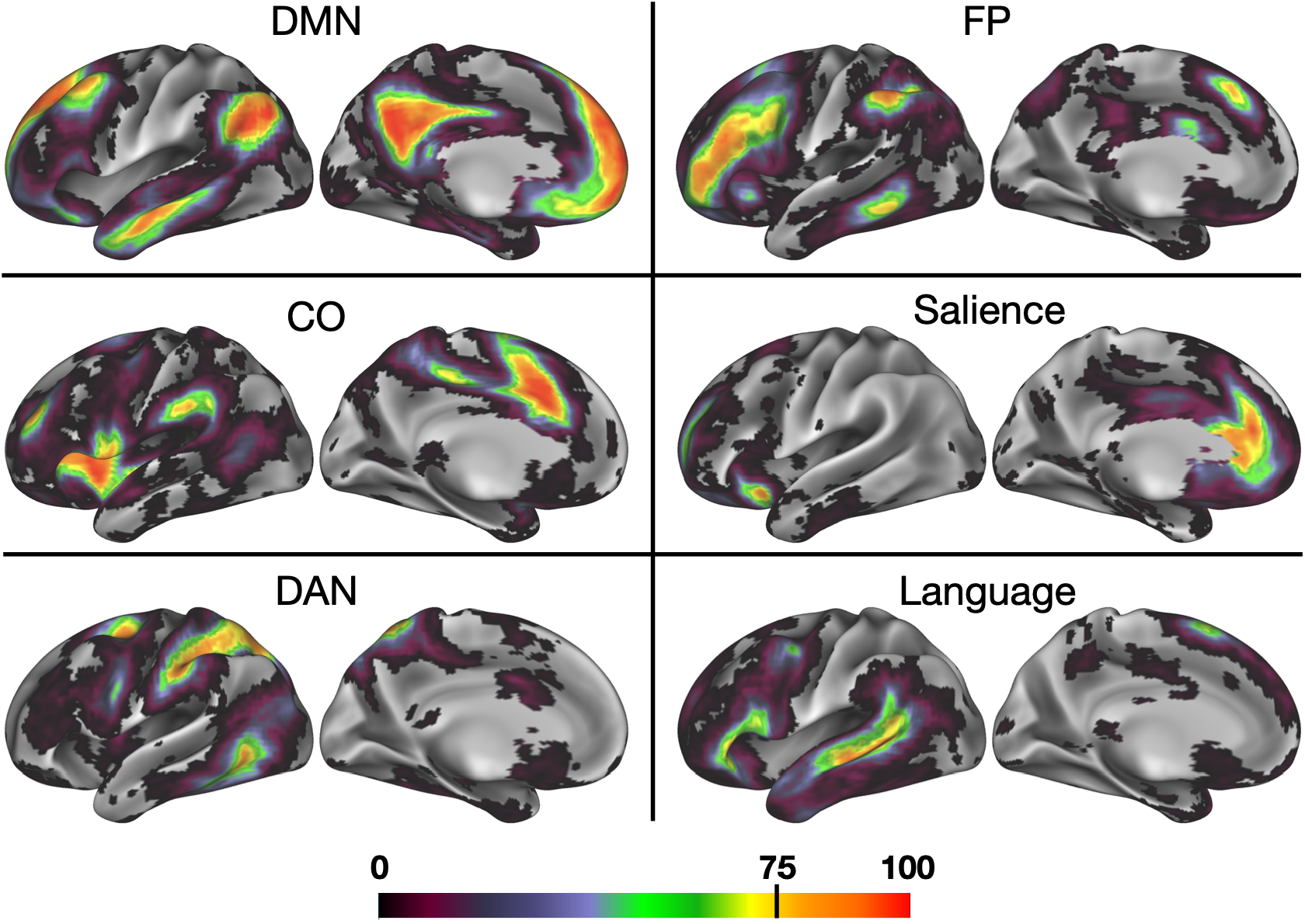
A probabilistic representation of 6 association networks. Cooler colors represent regions with the least confidence in network assignment across subjects, while warmer colors represent brain regions with the highest group consistency– in bright red regions, up to 100% of subjects converged on a given network assignment. (See Supp. Fig. 2 for probabilistic maps produced for both hemispheres and for all 14 networks.)

#### Consensus locations replicate across multiple datasets

Next, we implemented the probabilistic map procedure in three supplemental datasets (consisting of 9 MSC subjects, 384 HCP subjects, and 65 Yale subjects; Fig. 3). Despite differences in participant populations, scanners, and acquisition parameters (most notably in the HCP dataset), probabilistic network assignments generally replicate, with results from the three test datasets visually appearing similar at the 50 percent probability threshold and experiencing similar patterns of network “dropout” as the probability of assignment increases at 70 and 90 percent. We note that more dropout is observed in the HCP dataset, perhaps due to the lower SNR associated with these scans (e.g., see Seitzman et al., 2020, SI Fig. 4).

**Figure 3:**
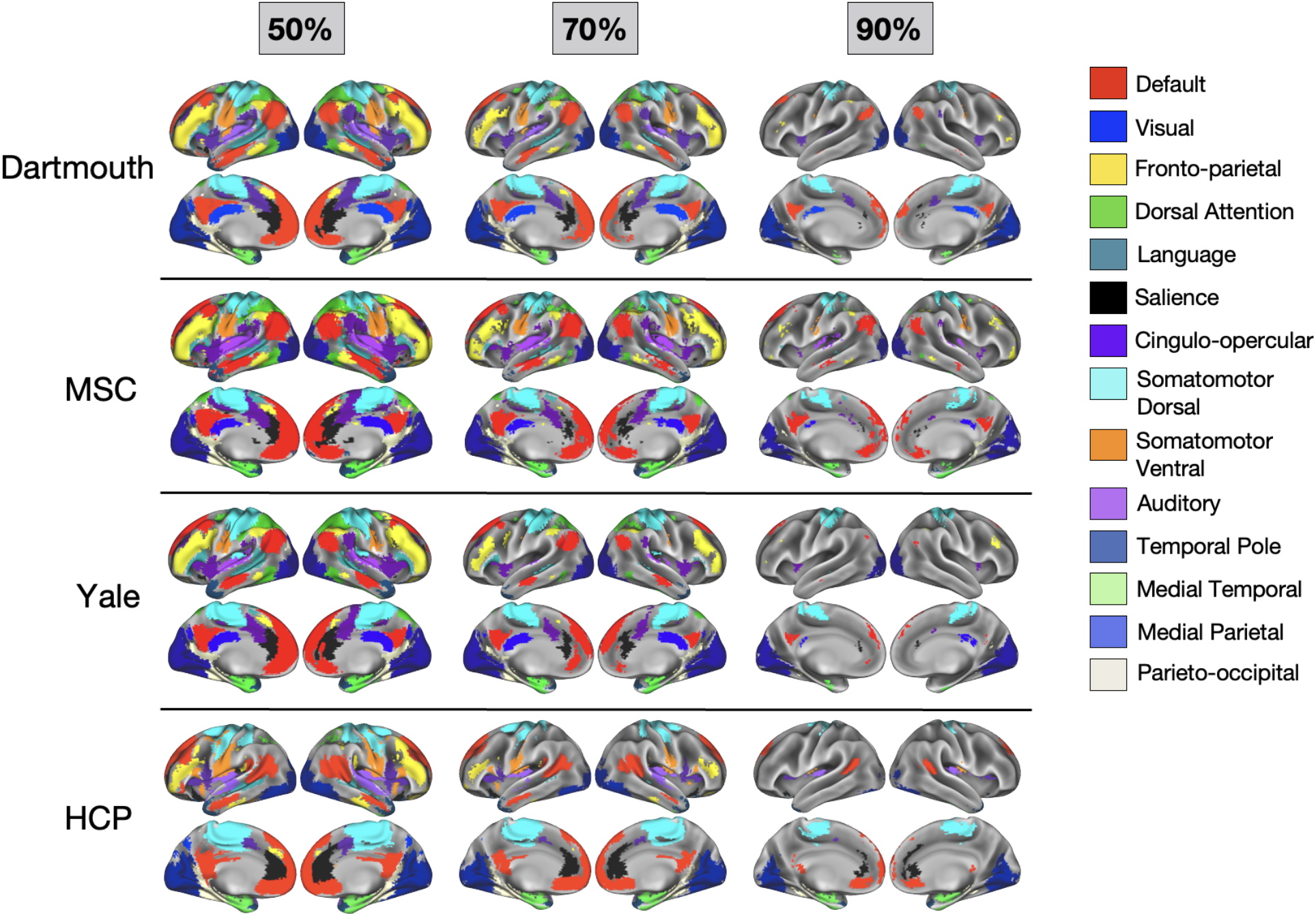
Thresholded probabilistic maps across 4 datasets. Probabilistic maps were generated for the primary Dartmouth datasets and from three additional datasets (MSC, HCP, and Yale). For each dataset, network assignments consistent across 50%, 70%, and 90% of subjects are displayed.

This observation was supported by quantitative comparisons as well. All three supplemental datasets showed a high spatial correlation with the Dartmouth dataset: on average, the network-specific probabilistic maps were correlated at r = 0.90 for Dartmouth:MSC, r = 0.90 for Dartmouth:Yale, and r = 0.70 for Dartmouth:HCP (see Supp. Fig. 4A for full breakdown by network). Furthermore, the high consensus locations (>70%) also replicated, as shown by network-wise rotation-based permutation analysis (p<0.001 relative to random null for all networks between Dartmouth and MSC and between Dartmouth and Yale; in the Dartmouth– HCP comparison, 10 of the 14 networks showed a Dice coefficient significantly higher than the null; see Supp. Fig. 4B for full break down by network).

Finally, in the highly sampled MSC dataset, we also demonstrate that probabilistic maps based on data-driven network assignments show high correspondence to the template-based assignments used here (Supp. Fig. 5). This suggests that template-matching and data-driven procedures converge with sufficient high-quality data, at least in neurotypical populations.

#### Individual networks vary in the size of their core and the span of surrounding components

While core regions of high consensus exist in all of the canonical networks investigated here, the networks vary in the extent of their more peripheral (i.e., low consensus) regions. As shown in Fig. 4, networks retain cortical territory at varying rates as the probability threshold (i.e., consensus across subjects) increases. For example, while the visual network consistently remains the most highly represented network across the highest probability thresholds, the inverse is true for FP: it is the third-most highly represented network across at least 50% of individuals, but when group consensus is examined at 80% or 90% of individuals, cortical representation of FP diminishes significantly.

**Figure 4:**
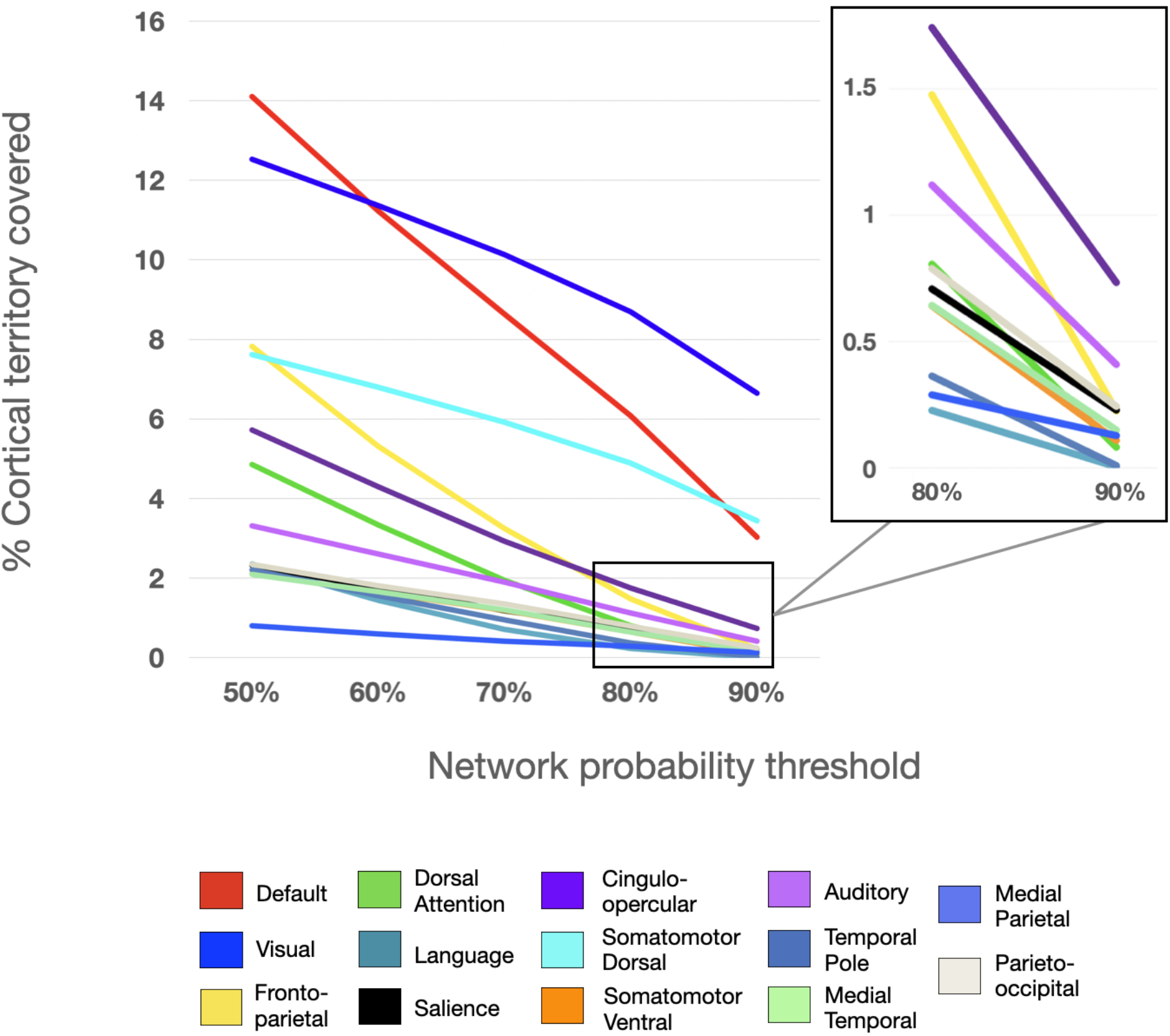
Representation of the proportion of cortical territory covered by each network at each probability threshold. Each line represents the total percentage of the cortex covered at a given threshold. Inset shows percent of cortical territory for the smaller networks at 80-90% consensus.

Differences in the rate of network “dropout” seem not to be driven purely by a distinction of sensorimotor vs. association networks. While sensorimotor networks tend to have higher consensus, some association networks also maintain a relatively high group consensus across thresholds, including DMN and CO. It appears unlikely that network size alone is driving the effect (i.e., that smaller networks taper off more quickly across probability thresholds); while some smaller networks experience relatively fast dropout (e.g., Lang.), others (e.g., PON and MTL) remain consistent across a high percentage of subjects. Regardless, all networks have some core regions of high inter-subject consensus, and networks vary in the cross-subject variability observed in locations surrounding these core regions.

#### Non-core areas overlap with previously described locations of network variants

Next, we sought to provide support for our approach by examining how consensus regions from this template-matching probabilistic procedure compared with previously identified locations of individual variability in functional network organization. Transparent white regions in Fig. 5 show “network variant” locations across 752 HCP subjects from Seitzman et al. (2019) where a given individual’s correlation pattern differs significantly from the group-average (this was computed using continuous measures, without reference to a discrete network assignment). Despite differences in methodological approaches used for identifying consensus probabilistic assignments (via template-matching) and individual variants (via low spatial correlations), we find that there is a good contrasting correspondence between these two methods. As would be expected, regions of high consensus lie mostly outside of the boundaries of network variants, and appear to fill in gaps where there is the greatest inter-subject variability in network assignment (e.g., temporoparietal junction, lateral frontal cortex).

**Figure 5:**
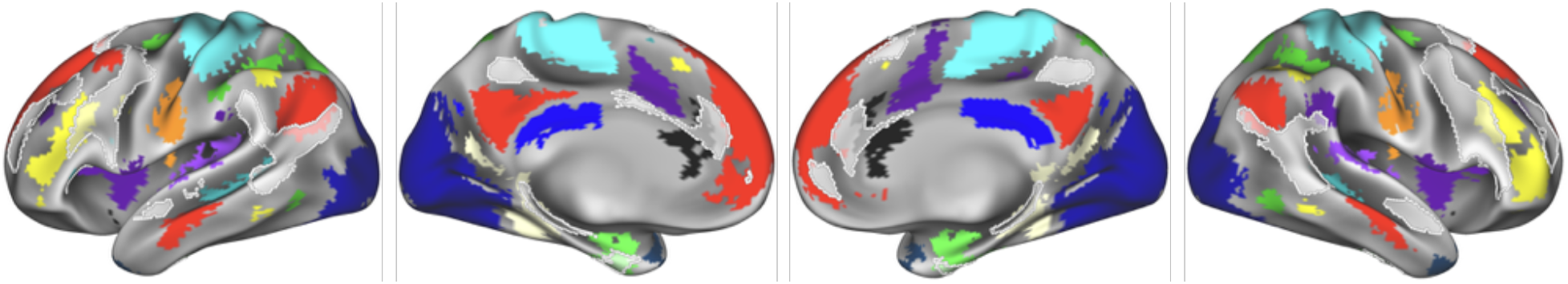
The spatial distribution of network variants across 752 HCP subjects (as identified in Seitzman et al., 2019) is displayed in transparent white, overlaid on the network map at 75% probability. The distribution displayed here is thresholded to show variant locations exhibited by at least 11% of subjects. Notably, the variants distribution appears to fill in gaps where there is the most inter-subject variability in network assignment, including temporoparietal junction and the left and right lateral frontal cortex.

#### Generation of a high-probability set of ROIs and point-and-click tool

A major goal of the current work was to improve group-studies by allowing researchers to evaluate network probabilities across participants and focus on locations of consensus. To this end, we sought to refine previous group-average ROI definitions based on these probabilistic network assignments to generate a set of high-consensus ROIs for future research. We began with the 248 cortical ROIs from the commonly used 264 regions from Power et al. (2011). We then restricted this set to regions where the average network assignment probability was ≥ 75% within the 7 mm diameter ROI. This resulted in 153 cortical ROIs. Thirteen of the 14 canonical networks were represented (no ROIs were retained for the temporal pole network), although the quantity of high-probability ROIs varied by network (see Fig. 6A for locations and network descriptions of ROIs). While the regions cover much of the cortex, some higher-variability areas such as the temporo-parietal junction and the lateral frontal cortex are more sparsely represented, as expected (e.g., see Gordon et al., 2017b; Laumann et al., 2015; Mueller et al., 2013; Seitzman et al., 2019). ROIs with the highest peak probabilities were identified largely in dorsal somatomotor and visual regions, with relatively lower peaks in lateral frontal and orbitofrontal regions (Fig. 6B)

**Figure 6:**
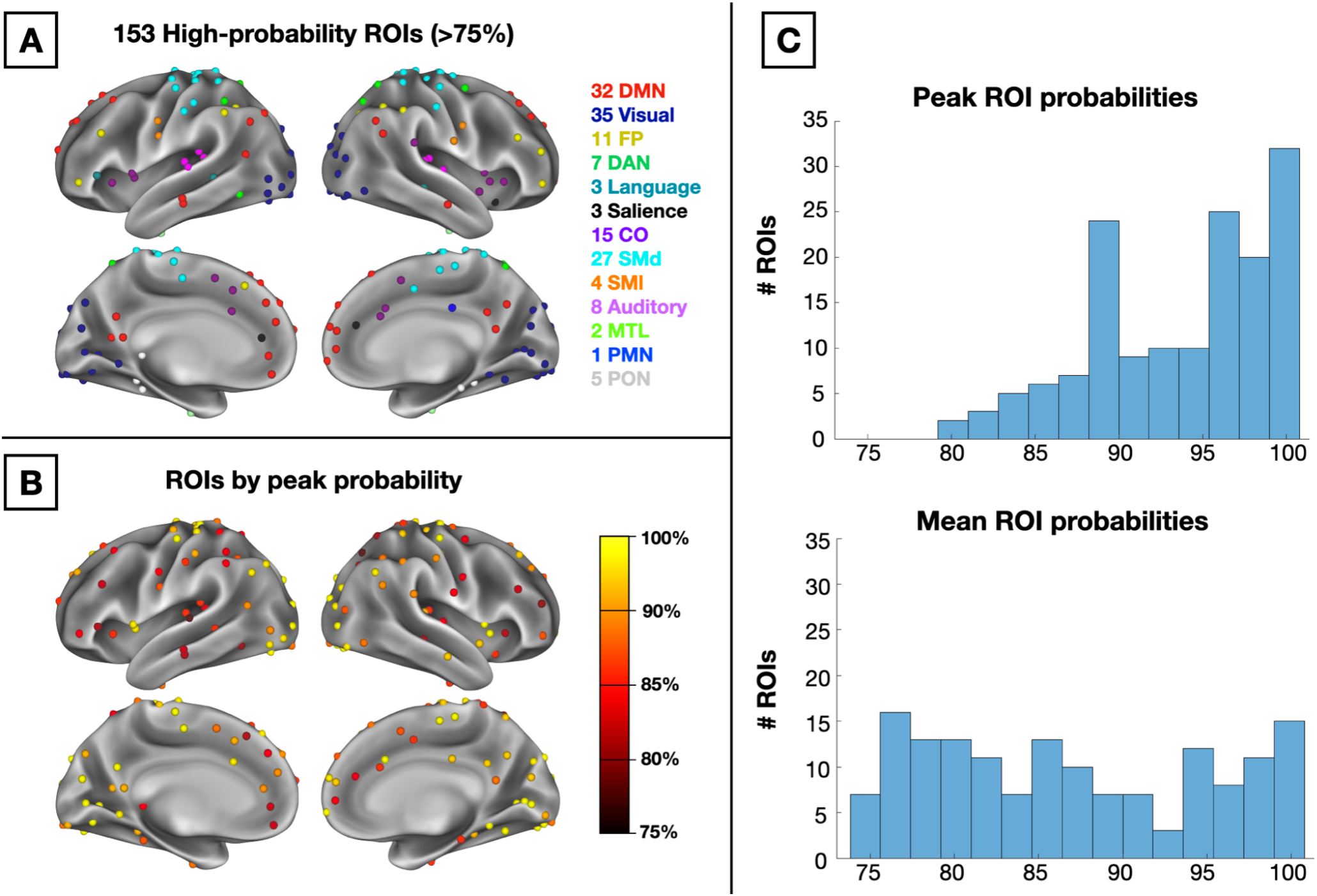
(A) 153 high-probability ROIs colored by their network description. (B) ROIs colored by peak probability across voxels within the ROI. (C) Histograms of peak probability values across all 153 ROIs (top) and mean probability values (bottom).

We provide each network’s probabilistic maps as a series of downloadable volume images, along with the 153 ROIs at https://github.com/GrattonLab (given the added differences seen with the HCP analyses, HCP-specific probabilistic maps are also provided). For researchers using Connectome Workbench, a *scene* file was created to allow researchers to explore network probabilities at every cortical voxel. Fig. 7 displays an example of this tool’s utility by exploring the DMN map, including an “Information” window with probabilities listed across all networks.

**Figure 7:**
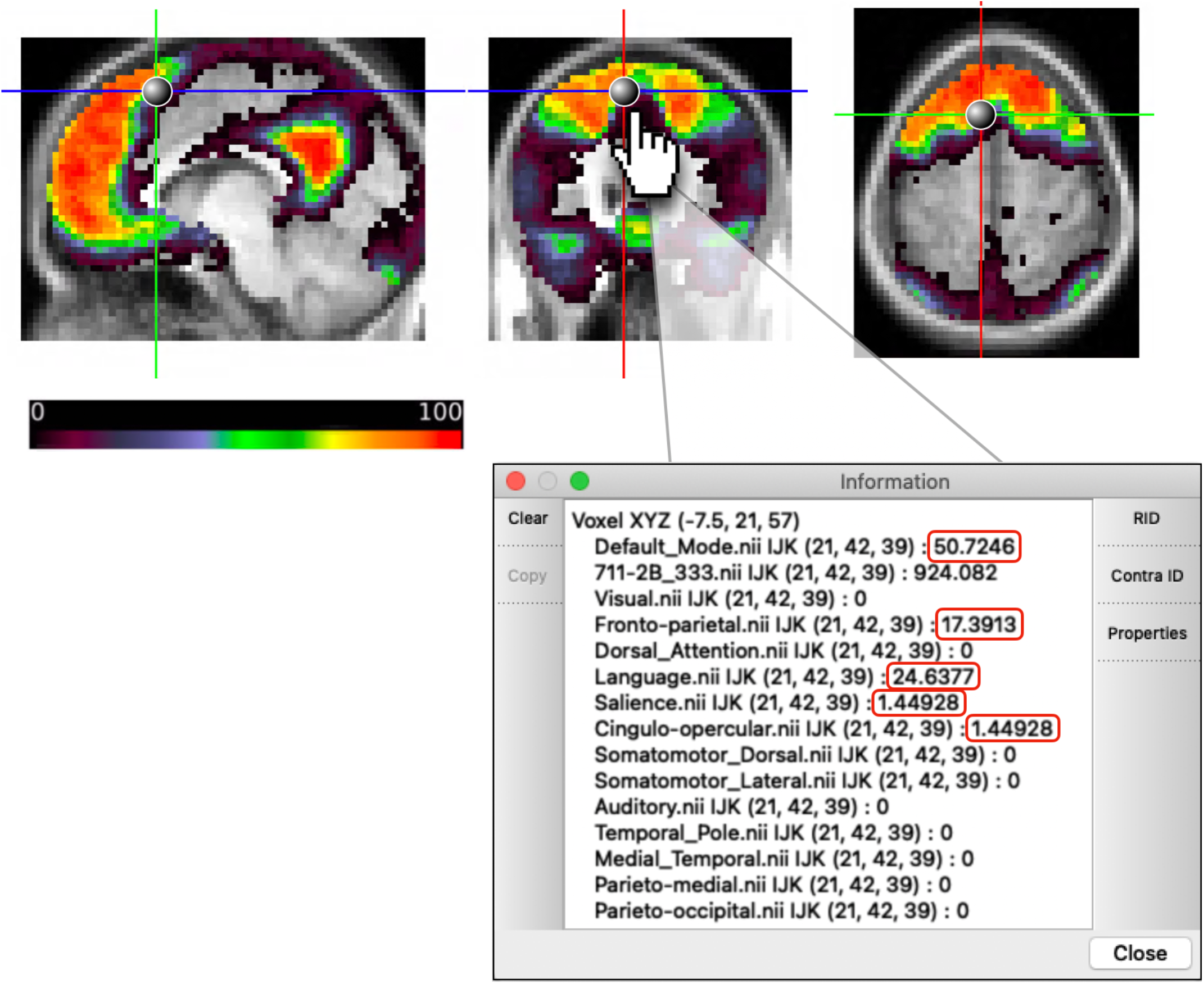
Schematic of publicly available research tool for exploring network probabilities. The DMN map is displayed, and probabilities of network membership to all 14 networks for the given voxel are listed in the “Information” window with non-zero probabilities outlined in red.

## DISCUSSION

Here, we probabilistically mapped functional networks across a group of highly sampled individuals. We found that there are “core” locations of high group consensus within each network. Networks vary in the extent and peak probability of their core regions, suggesting that networks with a higher group consensus may be more amenable to group-level analyses. The ability to identify locations with high group consensus allows for better-informed group studies of functional network properties, using either task or functional connectivity approaches. To facilitate this process, we provide a set of voxelwise probability maps for each of 14 canonical networks. In addition, we provide two tools for research use: (1) a set of network-specific, high-probability ROIs for use in task- and functional connectivity-based analyses and (2) a point-and-click tool allowing researchers to explore voxel-by-voxel probabilistic network estimates for regions of activation in their own data.

### Probabilistic approaches in imaging

In the imaging literature, probabilistic atlases have been utilized as a way to quantify spatial distributions of anatomical structures or functional areas to pinpoint locations of high consensus across a group. Many popular probabilistic atlases of the brain are based on anatomical data – e.g., the cerebellum (Diedrichsen et al., 2009), subcortical nuclei (Pauli et al., 2018), the basal ganglia (Keuken & Forstmann, 2015), tissue type, lobes, and sulci (Mazziotta et al., 1995) – to provide references for cross-subject comparisons. However, functional areas (at least in the cortex) do not necessarily conform well to anatomical definitions (Eickhoff et al., 2018; Gordon et al., 2016), suggesting that anatomical atlases are less well-suited for definition of functional ROIs in task-based or resting-state fMRI. The current cortical probabilistic atlas based on functional network mapping fills this gap. Future work in other age groups (e.g., youth, older adults) and clinical populations (e.g., schizophrenia, depression) may use a probabilistic approach to build additional probabilistic maps of functional networks and further enhance group studies in these domains.

### Utility of a probabilistic mapping approach to functional networks

We have adopted a probabilistic approach in this study given past evidence for both individual differences and group consensus in functional neuroanatomy (Gordon et al., 2017a; Gordon et al., 2017b; Gratton et al., 2018; Power et al., 2011; Yeo et al., 2011). It became increasingly apparent in our own work that, rather than qualitative statements about the magnitude of variability or the extent of similarity, of networks across individuals it would be useful to have a quantitative probabilistic view of the variability associated with each cortical location and each network to evaluate the consistency of our findings. Identifying regions of group consensus provides a wealth of opportunity for more well-informed research on brain networks.

The network probabilistic maps that we have produced can be used by researchers in a number of ways. First, these maps can be thresholded to create regions of interest for future (or retrospective) group analyses. For example, these maps may be thresholded to select regions of the frontoparietal and cingulo-opercular systems where we have high confidence in group consensus. This would allow for a re-analysis of past task dissociations between these two systems (Dubis et al., 2016; Gratton et al., 2017; Neta et al., 2015; Power & Petersen, 2013), but now accounting for potential individual variability in network assignments. To facilitate this application, we have provided a set of 153 ROIs that identify high-consensus regions within 13 of the 14 networks examined. Should a researcher wish to perform task-based or rest-based analyses at the group level, high-probability ROIs would be crucial in ensuring that the brain regions being analyzed are those which are most consistent across individuals; researchers can be more assured that a majority of individuals are providing information from the same network.

Secondly, these regions may be used to help interpret ambiguous results in group studies. For instance, a region which is assigned to the CO network in 70% of subjects and the salience network in the other 30% of subjects may serve as a meaningful distinction from a region which is assigned to CO in 70% of subjects but FP in the other 30%. Thus, while the high-probability ROIs focus on regions of group similarity, useful information on the locations and forms of individual variability can also be gleaned from the point and click probabilistic tool. In the future, researchers may use the probabilities associated with this paper to provide quantitative estimates for the typical (and atypical) network assignments associated with findings of interest.

Third, probabilistic network mapping may deepen our understanding of the clinical utility of mapping functional brain networks by providing reliable quantitative priors about the network assignments of each region. This probabilistic approach may provide a basis for more precisely identifying network deviations in individuals with specific diagnoses, as well as network changes across development. For example, one possible future investigation may be to examine whether individuals with a given clinical diagnosis vary predictably from the probability map of any network of interest; perhaps in clinical groups there will be more variability in higher-probability regions.

### Group consensus in core regions within large-scale networks

Our probabilistic maps demonstrated that each network was comprised of a set of “core” regions exhibiting very little or, in some cases, no variability (*note that our use of “core” is based on anatomical location, separate from the graph theoretical connotation of the word*). This suggests that the core areas of each network are relatively fixed across individuals, with little possibility for variation, and these regions complement previously described locations of high individual variability (see Fig. 5). The consensus areas of each network were larger in sensorimotor than association systems, consistent with the idea that association systems are more variable across individuals, maturation, and evolution, which has been suggested to be due to a lack of genetically encoded tethering markers in these areas (Buckner & Krienen, 2013). However, we found a consistent core in each of the association systems as well, which would appear to be at odds with a strong interpretation that association networks lack fixed constraints (Buckner & Krienen, 2013). Indeed, the consistency of association networks differed markedly between systems with, e.g., relatively robust consensus in the DMN and CO and high variability in the FP, despite their similar overall sizes and complex “high level” natures. Exploring the basis for commonalities and plasticity in association networks will be an interesting avenue for future work.

Importantly, while our results speak to areas of high and low variability in network assignment across subjects, less light is shed on locations that assign to multiple networks within a subject. The implementation of a template-matching approach to map networks in individuals, which necessarily forces a discrete network assignment, is not best suited to capture locations that may have network profiles intermediate to multiple networks. Some interpretations have characterized these regions as hubs (Gordon et al., 2018; Gratton et al., 2012, 2018; Power et al., 2013; Warren et al., 2014) while others describe them as multi-network integration zones reflective of a set of cortical gradients (Huntenburg et al., 2018). It will be interesting in future work to determine the correspondence between these “hub”-like intermediate regions that show inconsistent discrete network labels *within* a person and those that are variable *across* individuals. We note that the agreement between the current template-matching work and previous findings of individual differences based on continuous metrics (Seitzman et al., 2019) provides tentative evidence that these intermediate zones are not large contributors to the cross-person variability observed here.

### Limitations

The findings presented here have several limitations that are worth noting. First, in an effort to optimize the tradeoff between data quantity and the number of subjects retained for our probabilistic estimates, the amount of data required per subject was set to a minimum of 20 minutes of low-motion data. While this represents relatively higher-data subjects than a majority of group studies (which collect 5-10 min. of data), most of these subjects did not reach the 30-45 minute threshold that is ideal to produce asymptotic individual-subject reliability (Laumann et al., 2015). However, we were able to repeat the probabilistic analyses within the smaller but highly sampled Midnight Scan Club (MSC) dataset, which produced comparable results; in this dataset we were able to demonstrate that data-driven approaches show similar correspondence to the template-matching approaches used here (Supp. Fig. 5), further validating our findings.

Second, while there was general agreement in the probabilistic maps from the 4 datasets examined here, there were some differences, which may be driven by differences in scan parameters or dataset size/quality. This was particularly the case in the probabilistic map generated from the Human Connectome Project (HCP) dataset relative to the other three datasets. The probabilistic maps displayed in Fig. 3 reveal that some high-consensus regions that are conserved across probability thresholds in the Dartmouth, MSC, and Yale datasets show a lower degree of consensus in the HCP dataset. Such differences might be driven by the smaller voxel size and higher spatial and temporal resolution of the HCP dataset, which may lead to a lower signal-to-noise ratio (SNR; as demonstrated in Fig. S4 in Seitzman et al., 2020). Thus, the extent to which the probabilistic assignments replicate in datasets using similar acquisitions as the HCP is less certain and may require further investigation. Given this observation, we have also separately released the HCP-specific probabilistic network maps for use for those using the HCP dataset or others with similar acquisition parameters.

Lastly, we note that probabilistic assignments were calculated at the level of 14 canonical functional networks. This set of networks was selected because they are among those that have been most consistently defined and investigated in studies of cortical functional systems (Power et al., 2011; Yeo et al., 2011) and are thus likely to be useful to a broad set of individuals. However, this selection necessarily limits the observation of probabilistic maps at other resolutions, including those of interesting sub-network structure such as the default mode subnetworks identified by Braga & Buckner (2017), Gordon et al. (2020), and Kong et al. (2019). Moreover, the current approach is based on a probabilistic representation of the systems, not areal, level of brain organization. A consistent network assignment across individuals is not a guarantee that a region belongs to the same brain area across those individuals. We know from past work based on functional localizers that there is variability across subjects at the areal level as well (e.g., Kanwisher et al., 1997; Wang et al., 2015); variation at the areal level may also carry information about individual differences, and will be important in studies requiring area-level precision. An exciting avenue for future work is to expand on the techniques in this manuscript to probabilistically map sub-networks and areal level organization.

## CONCLUSIONS

Here, we produce a probabilistic representation of distributions of functional network assignments across a group of highly sampled subjects. While individual networks vary in the span of their “core” high-probability locations, all networks examined showed regions of high group consensus. These probabilistic maps and core regions replicated across four diverse datasets. The quantitative probabilistic maps, high-consensus ROIs, and point-and-click tool produced from these analyses will allow researchers to improve group studies by providing added information of cross-subject consensus.

## Acknowledgements

Funding was provided by NIH grant R01MH118370 (CG), the James S. McDonnell Foundation (SEP), and NIH grant R01MH111640 (MN).

## SUPPLEMENTARY MATERIALS

### Supplemental Methods

#### Template generation

Templates were generated based on a map of data-driven group-average network assignments in the cortex, using a related approach to Seitzman et al. (2019) and Gordon et al. (2017). These data-driven group-average network assignments were produced from the WashU-120 dataset, and the more highly sampled WashU-24 dataset was then used to generate a final high-quality network average (template).

After applying a cortical gray matter mask for each subject in the WashU-120, a voxelwise correlation matrix was calculated by correlating each cortical voxel’s BOLD timeseries with the timeseries of every other cortical voxel, resulting in a 57,544 by 57,544 matrix. Correlation matrices were Fisher transformed and averaged across subjects. The inverse Fisher transform was then applied to the resulting group-average matrix.

Next, the data-driven community detection InfoMap algorithm (Rosvall & Bergstrom, 2008) was applied to the group average matrix to identify cortical brain networks. Network definition in the group-average involved computing community assignments at a range of thresholds (0.5% to 5% by increments of 0.1%, similar to Gordon et al., 2017). Voxel-wise connections within 20 mm of each other were excluded from consideration in order to avoid correlations to be dominated by the effects of spatial smoothing, following Laumann et al. (2015) and Gordon et al. (2017).

A consensus assignment was then derived by an automated algorithm (Seitzman et al., 2020), which summarizes assignments by weighting communities across thresholds. The algorithm allows smaller networks to contribute more heavily by allotting greater weight to the sparser thresholds at which those communities are commonly detected. This procedure resulted in 14 networks, which are consistent with those identified by previous versions in Gordon et al. (2017) and Laumann et al. (2015) based on a surface-processing stream.

Next, templates were created for each of these data-driven networks using the more highly sampled WashU-24 in an effort to provide the highest quality templates possible. The timeseries for each network was extracted for each subject (averaged across voxels assigned to that network). This average timeseries was used to create a network seedmap by correlating the timeseries will all other gray matter voxels; this resultant map was averaged across the WashU-24 subjects. This procedure was repeated for each network, producing a group-average template connectivity map for each brain network (see Supp. Fig. 1 for a schematic of the creation of network templates and a view of all network templates).

## Supplemental Tables

**Supplementary Table 1**

**Supp. Table 1.** Acquisition parameters for functional MRI runs.

| Dataset | Scanner | TR (s) | TE (ms) | Flip angle | Num. slices | Voxel size (mm) |
| --- | --- | --- | --- | --- | --- | --- |
| WashU-120 and WashU-24 | Siemens Trio 3T | 2.5 | 27 | 90° | 32-36 | 4 x 4 x 4 |
| Dartmouth | Philips Achieva 3T | 2.5 | 35 | 90° | 36 | 3 x 3 x 3.5 |
| MSC | Siemens Trio 3T | 2.2 | 27 | 90° | 36 | 4 x 4 x 4 |
| HCP | Custom HCP Skyra 3T | 0.72;<br>MB = 8 | 33 | 52° | 72 | 2 x 2 x 2 |
| Yale | Siemens Trio 3T | 1.55 | 30 | 80° | 25 | 3.4 x 3.4 x 6 |

## Supplemental Figures

**Supplementary Fig. 1**

**Supp. Fig. 1.**
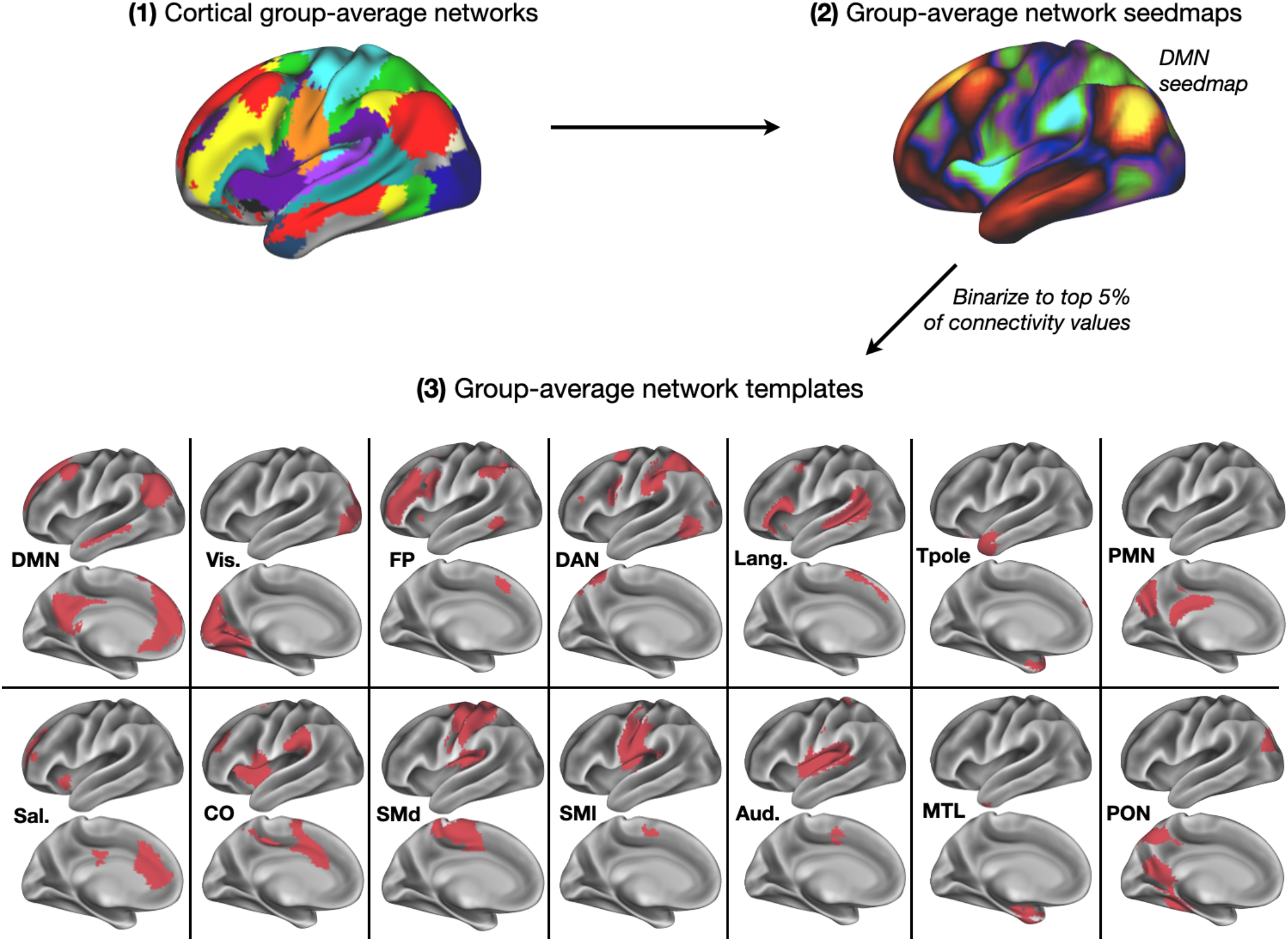
Creation of network templates and binarized versions of network templates for each of the 14 networks analyzed in this study.

**Supplementary Fig. 2**

**Supp. Fig. 2.**
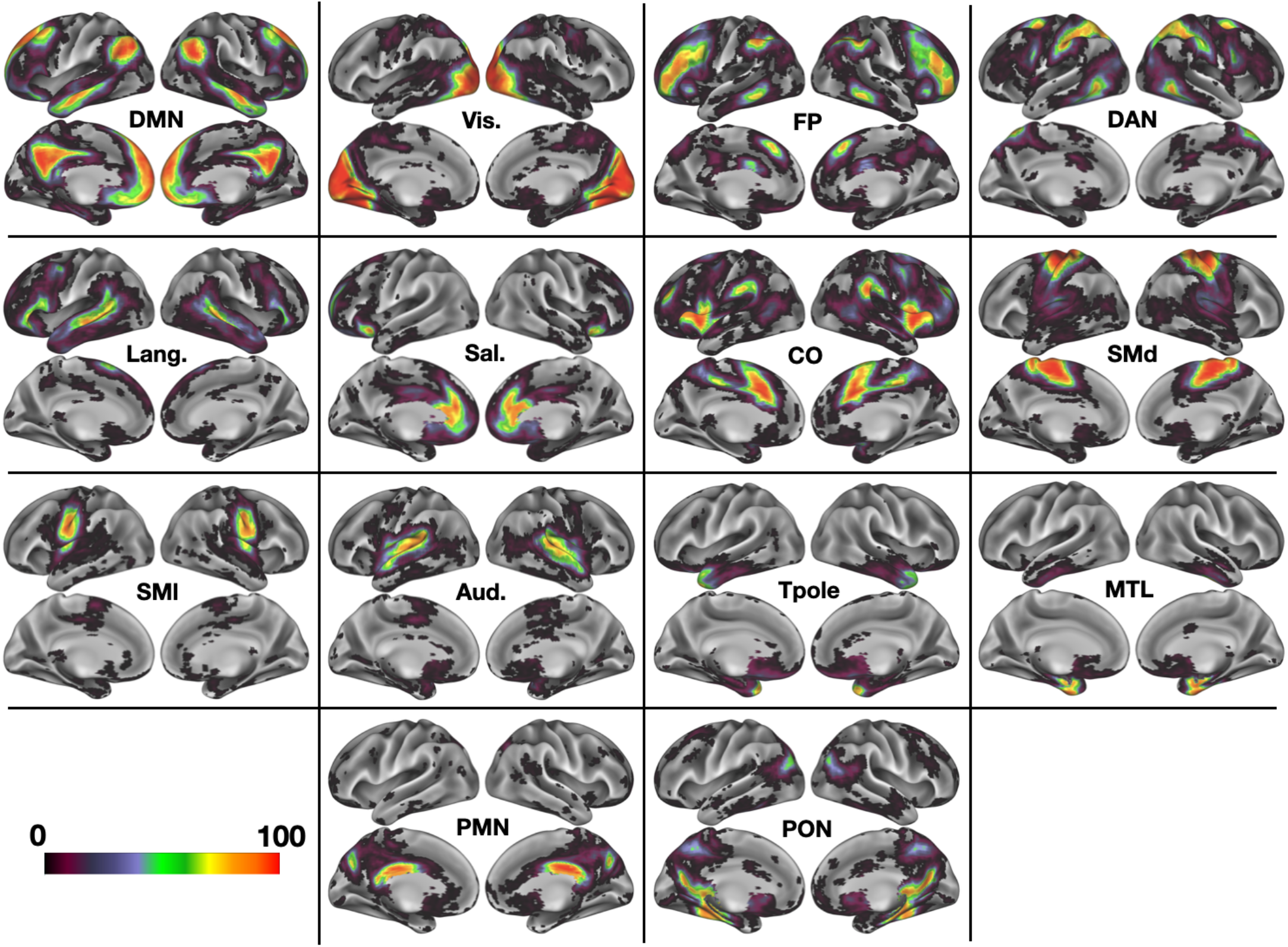
Probabilistic overlap maps for all 14 networks.

**Supplementary Fig. 3**

**Supp. Fig. 3.**
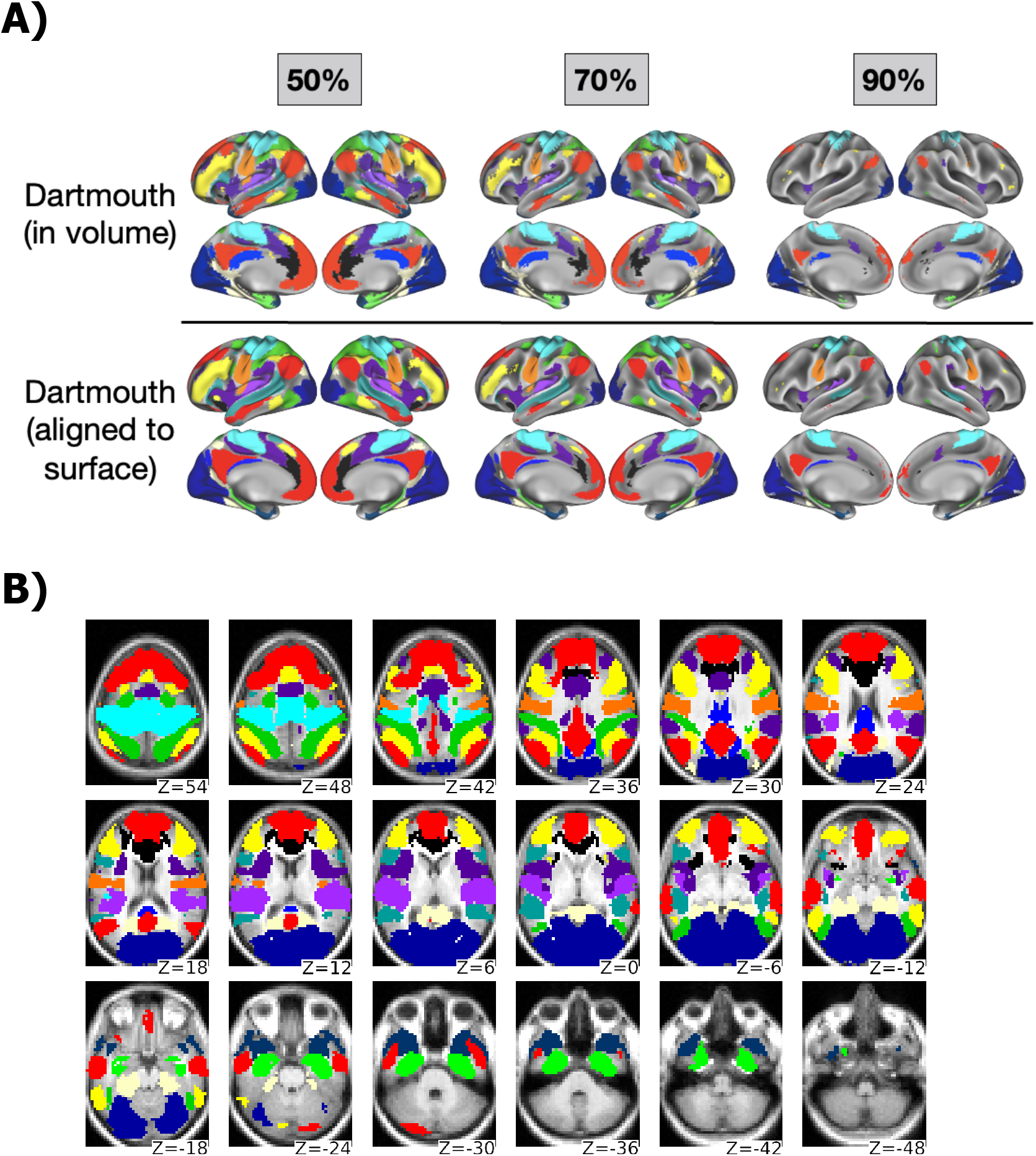
(A) Comparison of probabilistic network estimates based on volume- vs. surface-based alignment. For the surface-based alignment, surfaces were generated with FreeSurfer’s recon-all pipeline, native surfaces were aligned to the fsaverage surface by shape-based spherical registration, and BOLD timeseries were mapped to the surface following procedures outlined in (Gordon et al., 2016). Probabilistic outcomes are extremely similar across processing methods and do not appear to be greatly impacted by such methodological differences. (B) Thresholded probabilistic network maps at 50% probability are displayed in the original volume space where processing was performed to show the effects of smoothness in the volume.

**Supplementary Fig. 4**

**Supp. Fig. 4.**
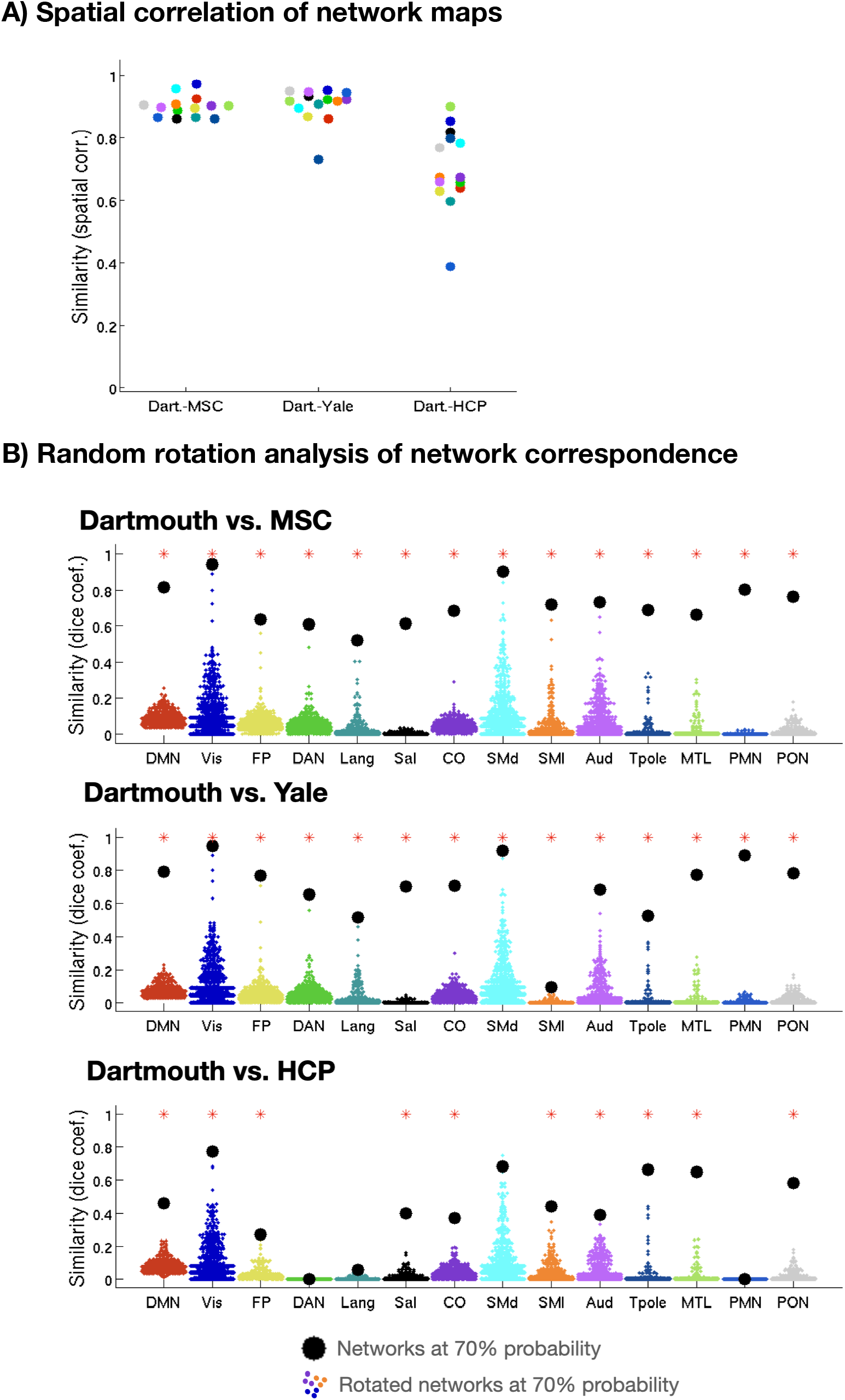
(A) Spatial correlation of each network’s continuous (unthresholded) probabilistic map between the primary Dartmouth dataset and three others (MSC, Yale, and HCP). Average *r* for Dartmouth–MSC = 0.90; average for Dartmouth–Yale = 0.90; average for Dartmouth–HCP = 0.70. (B) Overlap of high consensus (>70%) locations was quantified using a rotation-based analysis, which compared randomly rotated surface-mapped networks in the Dartmouth dataset to the replication datasets via the Dice coefficient. Average Dice coefficients (as calculated across networks) were 0.72 for the Dartmouth–MSC comparison, 0.70 for Dartmouth–Yale, and 0.41 for Dartmouth–HCP.

**Supplementary Fig. 5**

**Supp. Fig. 5.**
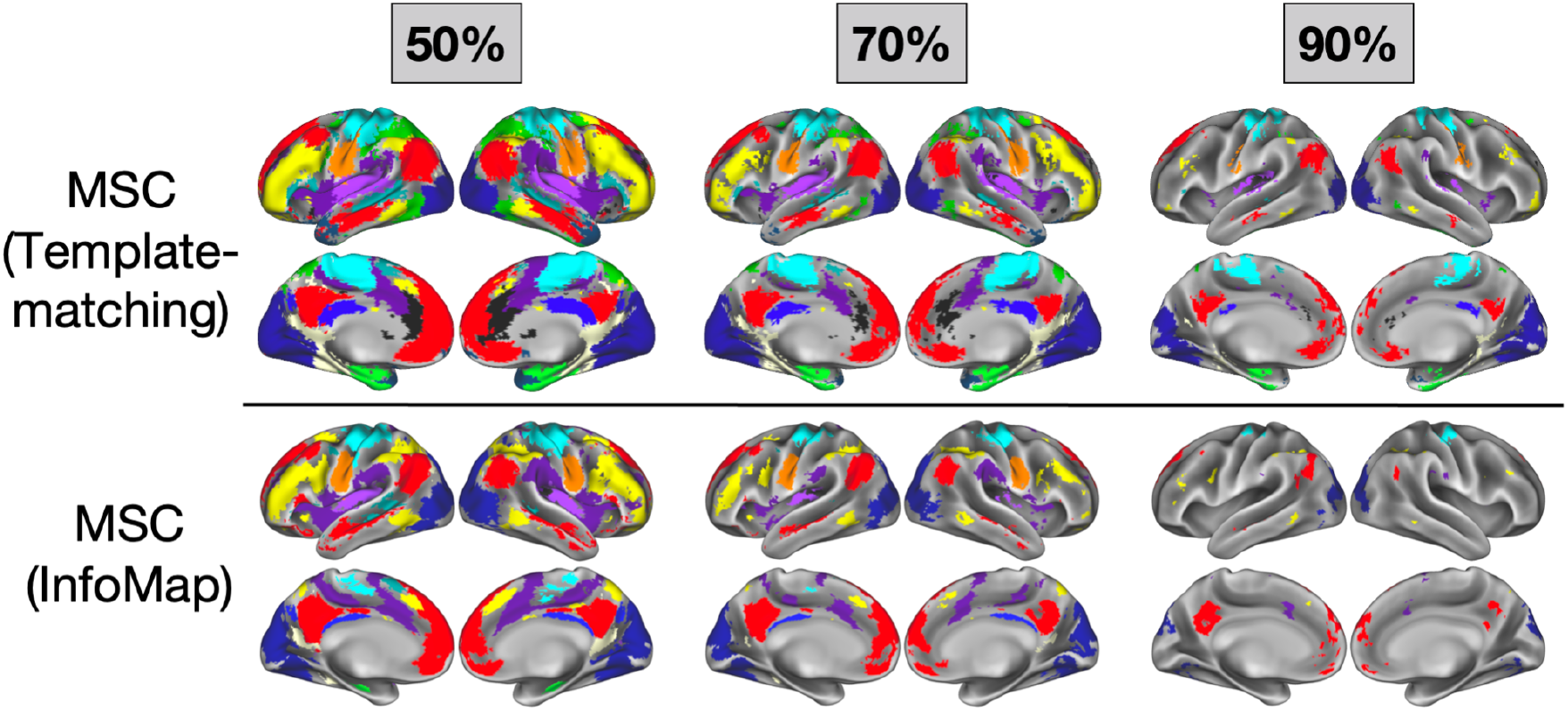
The use of a data-driven technique (InfoMap) to derive network assignments in the highly sampled MSC dataset produces probabilistic maps that shows good correspondence to the template-matching approach used here, despite the lower level of constraints in this technique.

